## Supplemental information for "Molecular evolution of CO_2_-sensing ab1C neurons underlies divergent sensory responses in the *Drosophila suzukii* species group"

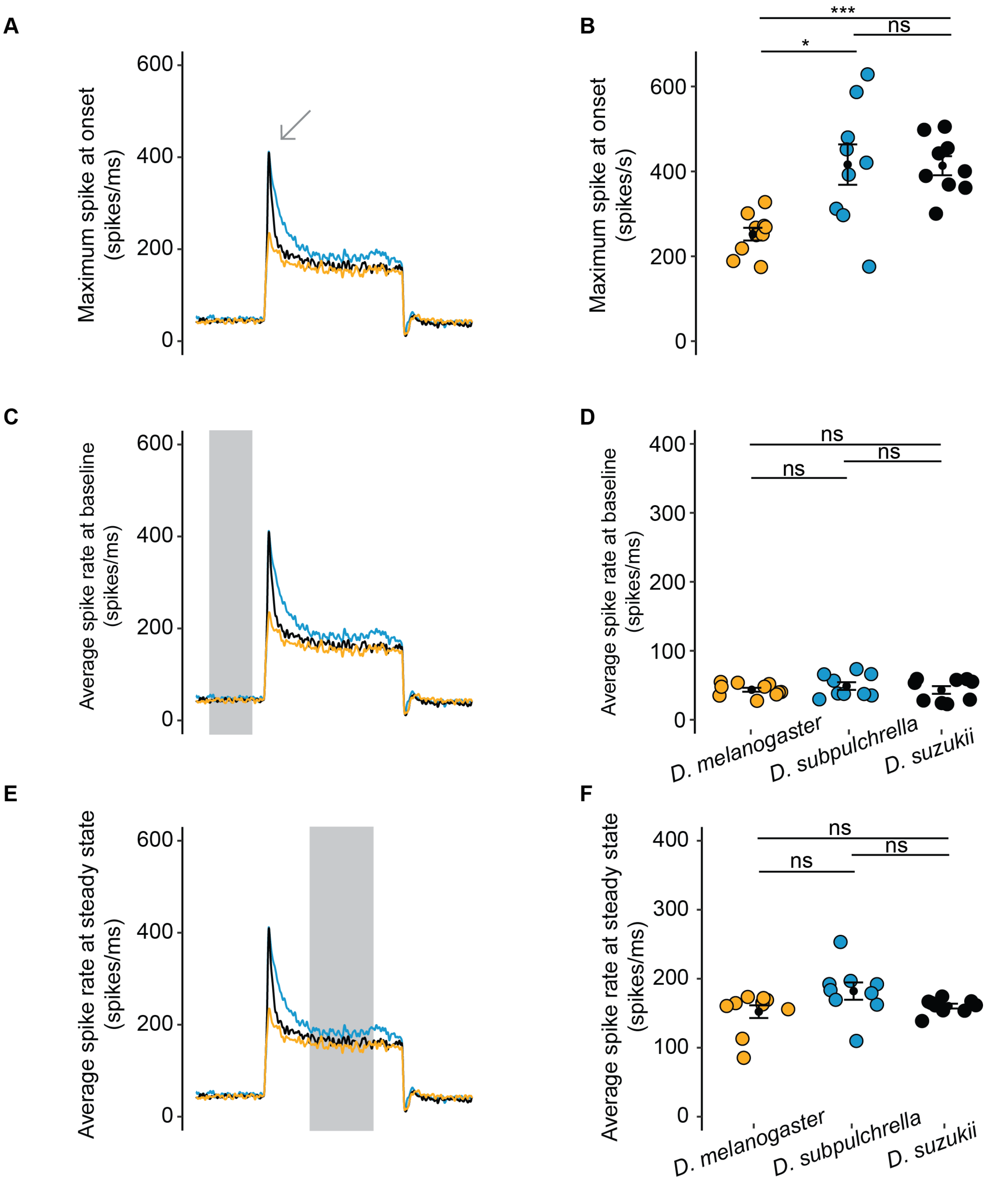


S1 Fig: Single sensillum electrophysiology recordings for the ab1c neuron in wildtype flies.
A-B) The maximum spike rate across each ab1c neuron recording is significantly higher in *D. suzukii* and *D. subpulchrella* compared to *D. melanogaster* (Kruskal-Wallis followed by pairwise Wilcoxon signed-rank test with Holm correction, *Q* = 0.000, Q = 0.017). C-D) The ab1c spike rate of the three species is not significantly different at baseline (Kruskal-Wallis followed by pairwise Wilcoxon signed-rank test with Holm correction, Dsub-Dmel: Q = 1.00, Dsuz-Dmel: Q = 1.00, Dsub-Dsuz: Q=0.89). E-F) The ab1c spike rate of the three species is not significantly different at steady state, which was calculated across

1000 ms (Kruskal-Wallis followed by pairwise Wilcoxon signed-rank test with Holm correction, Dsub-Dmel: Q = 0.066, Dsuz-Dmel: Q = 1.00, Dsub-Dsuz: Q= 0.056).

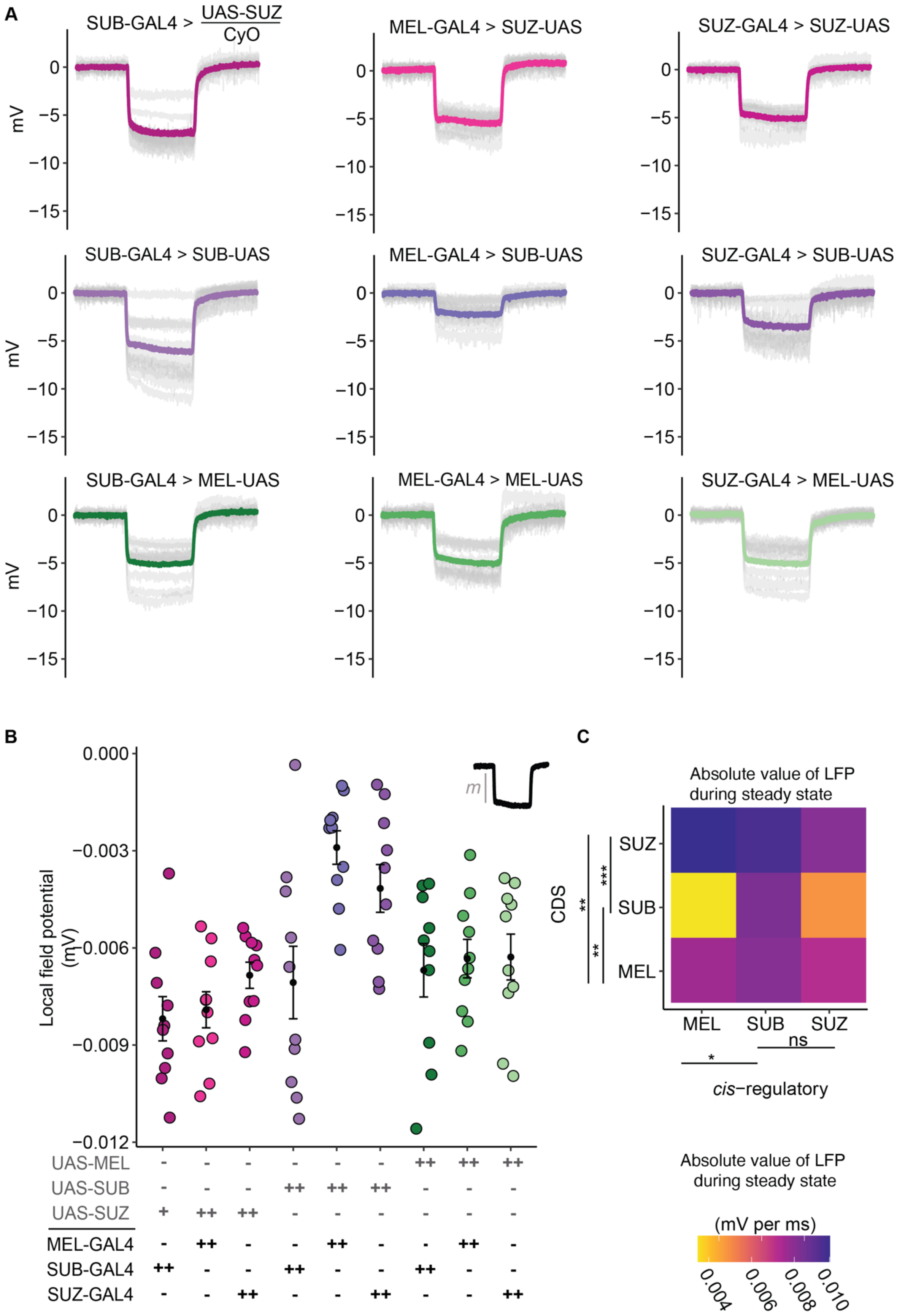

A
SUB-GAL4 > UAS-SUZ / CyO
MEL-GAL4 > SUZ-UAS
SUZ-GAL4 > SUZ-UAS
SUB-GAL4 > SUB-UAS
MEL-GAL4 > SUB-UAS
SUZ-GAL4 > SUB-UAS
SUB-GAL4 > MEL-UAS
MEL-GAL4 > MEL-UAS
SUZ-GAL4 > MEL-UAS
mV
0
−5
−10
−15
B
Local field potential (mV)
0.000
−0.003
−0.006
−0.009
−0.012
m
UAS-MEL
UAS-SUB
UAS-SUZ
MEL-GAL4
SUB-GAL4
SUZ-GAL4
C
Absolute value of LFP during steady state
CDS
SUZ
SUB
MEL
MEL SUB SUZ
cis−regulatory
ns
*
**
***
Absolute value of LFP during steady state
(mV per ms)
0.004
0.006
0.008
0.010

S2 Fig: Species-specific effects of Gr63a coding and regulatory variation on $CO_2$-evoked neural activity. A) LFP recordings for each transgenic line of the ab1c neurons recorded in Fig 4. Each grey line is the average of five replicates from one ab1c neuron, and the colored lines are the average across ten neurons. B) Without correcting for the heterozygous individual, the slope of the LFP at $CO_2$ onset was calculated across 750 ms. For the *CDS,* transgenic lines with *D. suzukii* CDS had the highest rate of change, followed by *D. melanogaster* and then *D. subpulchrella* (Kruskal-Wallis followed by pairwise Wilcoxon signed-rank test with Holm correction, Dsub-Dsuz: $Q = 0.00027$*,* Dsuz-Dmel: $Q = 0.01990$, Dsub-Dmel: $Q = 0.01836$). For the *cis*-regulatory element, the *D. subpuclrehlla* had the highest rate of change and trended lower than the other species elements (Kruskal-Wallis followed by pairwise Wilcoxon signed-rank test with Holm correction, Dsub-Dsuz: $Q = 0.092$*,* Dsuz-Dmel: $Q = 0.982$, Dsub-Dmel: $Q = 0.092$).

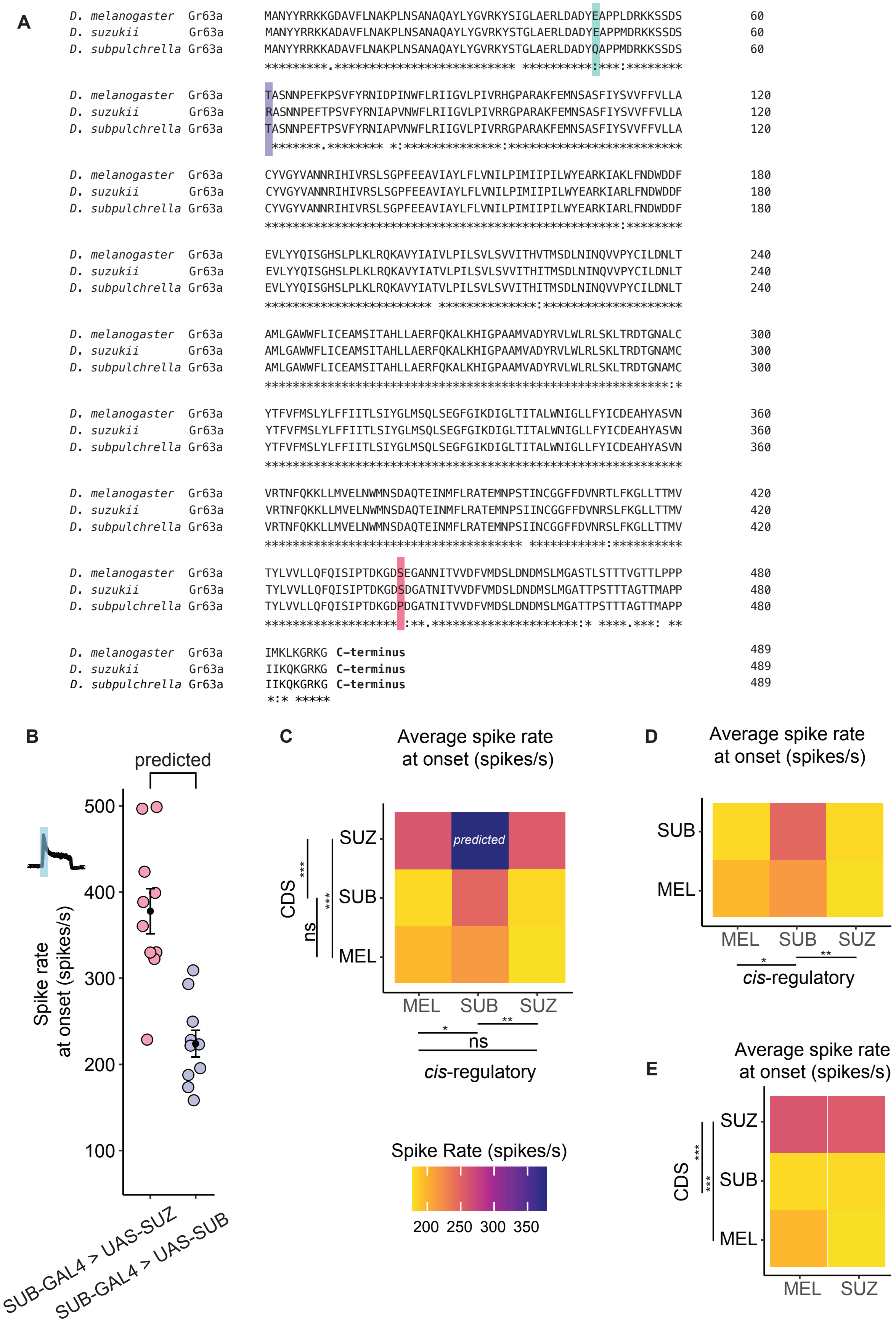

A
D. melanogaster Gr63a MANYYRRKKGDAVFLNAKPLNSANAQAYLYGVRKYSIGLAERLDADYEAPPLDRKKSSDS 60
D. suzukii Gr63a MANYYRRKKADAVFLNAKPLNSANAQAYLYGVRKYSTGLAERLDADYEAPPMDRKKSSDS 60
D. subpulchrella Gr63a MANYYRRKKADAVFLNAKPLNSANAQAYLYGVRKYSTGLAERLDADYQAPPMDRKKSSDS 60
*********.************************** **********:***:********
D. melanogaster Gr63a TASNNPEFKPSVFYRNIDPINWFLRIIGVLPIVRHGPARAKFEMNSASFIYSVVFFVLLA 120
D. suzukii Gr63a RASNNPEFTPSVFYRNIAPVNWFLRIIGVLPIVRRGPARAKFEMNSASFIYSVVFFVLLA 120
D. subpulchrella Gr63a TASNNPEFTPSVFYRNIAPVNWFLRIIGVLPIVRRGPARAKFEMNSASFIYSVVFFVLLA 120
*******.******** *:**************:************************
D. melanogaster Gr63a CYVGYVANNRIHIVRSLSGPFEEAVIAYLFLVNILPIMIIPILWYEARKIAKLFNDWDDF 180
D. suzukii Gr63a CYVGYVANNRIHIVRSLSGPFEEAVIAYLFLVNILPIMIIPILWYEARKIARLFNDWDDF 180
D. subpulchrella Gr63a CYVGYVANNRIHIVRSLSGPFEEAVIAYLFLVNILPIMIIPILWYEARKIARLFNDWDDF 180
****************************************************:********
D. melanogaster Gr63a EVLYYQISGHSLPLKLRQKAVYIAIVLPILSVLSVVITHVTMSDLNINQVVPYCILDNLT 240
D. suzukii Gr63a EVLYYQISGHSLPLKLRQKAVYIATVLPILSVLSVVITHITMSDLNINQVVPYCILDNLT 240
D. subpulchrella Gr63a EVLYYQISGHSLPLKLRQKAVYIATVLPILSVLSVVITHITMSDLNINQVVPYCILDNLT 240
************************ **************:********************
D. melanogaster Gr63a AMLGAWWFLICEAMSITAHLLAERFQKALKHIGPAAMVADYRVLWLRLSKLTRDTGNALC 300
D. suzukii Gr63a AMLGAWWFLICEAMSITAHLLAERFQKALKHIGPAAMVADYRVLWLRLSKLTRDTGNAMC 300
D. subpulchrella Gr63a AMLGAWWFLICEAMSITAHLLAERFQKALKHIGPAAMVADYRVLWLRLSKLTRDTGNAMC 300
**********************************************************:*
D. melanogaster Gr63a YTFVFMSLYLFFIITLSIYGLMSQLSEGFGIKDIGLTITALWNIGLLFYICDEAHYASVN 360
D. suzukii Gr63a YTFVFMSLYLFFIITLSIYGLMSQLSEGFGIKDIGLTITALWNIGLLFYICDEAHYASVN 360
D. subpulchrella Gr63a YTFVFMSLYLFFIITLSIYGLMSQLSEGFGIKDIGLTITALWNIGLLFYICDEAHYASVN 360
************************************************************
D. melanogaster Gr63a VRTNFQKKLLMVELNWMNSDAQTEINMFLRATEMNPSTINCGGFFDVNRTLFKGLLTTMV 420
D. suzukii Gr63a VRTNFQKKLLMVELNWMNSDAQTEINMFLRATEMNPSIINCGGFFDVNRSLFKGLLTTMV 420
D. subpulchrella Gr63a VRTNFQKKLLMVELNWMNSDAQTEINMFLRATEMNPSIINCGGFFDVNRSLFKGLLTTMV 420
************************************ ***********:**********
D. melanogaster Gr63a TYLVVLLQFQISIPTDKGDSEGANNITVVDFVMDSLDNDMSLMGASTLSTTTVGTTLPPP 480
D. suzukii Gr63a TYLVVLLQFQISIPTDKGDSDGATNITVVDFVMDSLDNDMSLMGATTPSTTTAGTTMAPP 480
D. subpulchrella Gr63a TYLVVLLQFQISIPTDKGDPDGATNITVVDFVMDSLDNDMSLMGATTPSTTTAGTTMAPP 480
******************* :**.********************:* ****.***: **
D. melanogaster Gr63a IMKLKGRKG C-terminus 489
D. suzukii Gr63a IIKQKGRKG C-terminus 489
D. subpulchrella Gr63a IIKQKGRKG C-terminus 489
*:* *****
B
predicted
Spike rate at onset (spikes/s)
500
400
300
200
100
SUB-GAL4 > UAS-SUZ
SUB-GAL4 > UAS-SUB
C
Average spike rate at onset (spikes/s)
SUZ
SUB
MEL
predicted
CDS
***
***
ns
MEL SUB SUZ
*
**
ns
cis-regulatory
Spike Rate (spikes/s)
200 250 300 350
D
Average spike rate at onset (spikes/s)
SUB
MEL
MEL SUB SUZ
*
**
cis-regulatory
E
Average spike rate at onset (spikes/s)
SUZ
SUB
MEL
CDS
***
***
MEL SUZ

S3 Fig. Comparison of *Gr63a* coding sequences and spike rates of transgenic lines. A) Comparison of *Gr63a* coding sequences among *D. suzukii*, *D. subpulchrella*, and *D. melanogaster.* B) Predicted spike rates at CO2 onset for the SUB-GAL4> UAS-SUZ individual as well as the SUB-GAL4 > UAS-SUB control. The predicted spike rate of SUB-GAL4 > SUB-UAS and the actual spike rate are very similar (Kruskal-Wallis followed by pairwise Wilcoxon signed-rank test with Holm correction, Q = 0.19, mean of B = 223.721, mean of estimated B = 228.3726), which gave us confidence that the predicted homozygous spike rate for SUB-GAL4 > UAS-SUZ is robust. C) Heat map comparing the spike rate between homozygous transgenic lines, including the modeled SUB-GAL4 >UAS-SUZ value, carrying *subpulchrella* and *melanogaster* proteins with *cis*-regulatory sequences from the three species. The *D. suzukii* CDS (Kruskal-Wallis followed by pairwise Wilcoxon signed-rank test with Holm correction, Dmel–Dsuz: Q = 1.1e-08 and Dsub-Dsuz: Q = 6.3e-07) and *D. subpulchrella* cis-regulatory element (Kruskal-Wallis followed by pairwise Wilcoxon signed-rank test with Holm correction, Q = 0.0139 between *D. melanogaster* and *D. subpulchrella* and Q = 0.004 between *D. subpulchrella* and *D. suzukii*) resulted in the highest spike rate. D) Spike rate comparisons of homozygote transgenic lines carrying *subpulchrella* and *suzukii* proteins with regulatory sequences from the three species. E) Spike rate comparisons of homozygote transgenic lines carrying *melanogaster* and *suzukii* regulatory sequences with CDS from the three species.

A

B

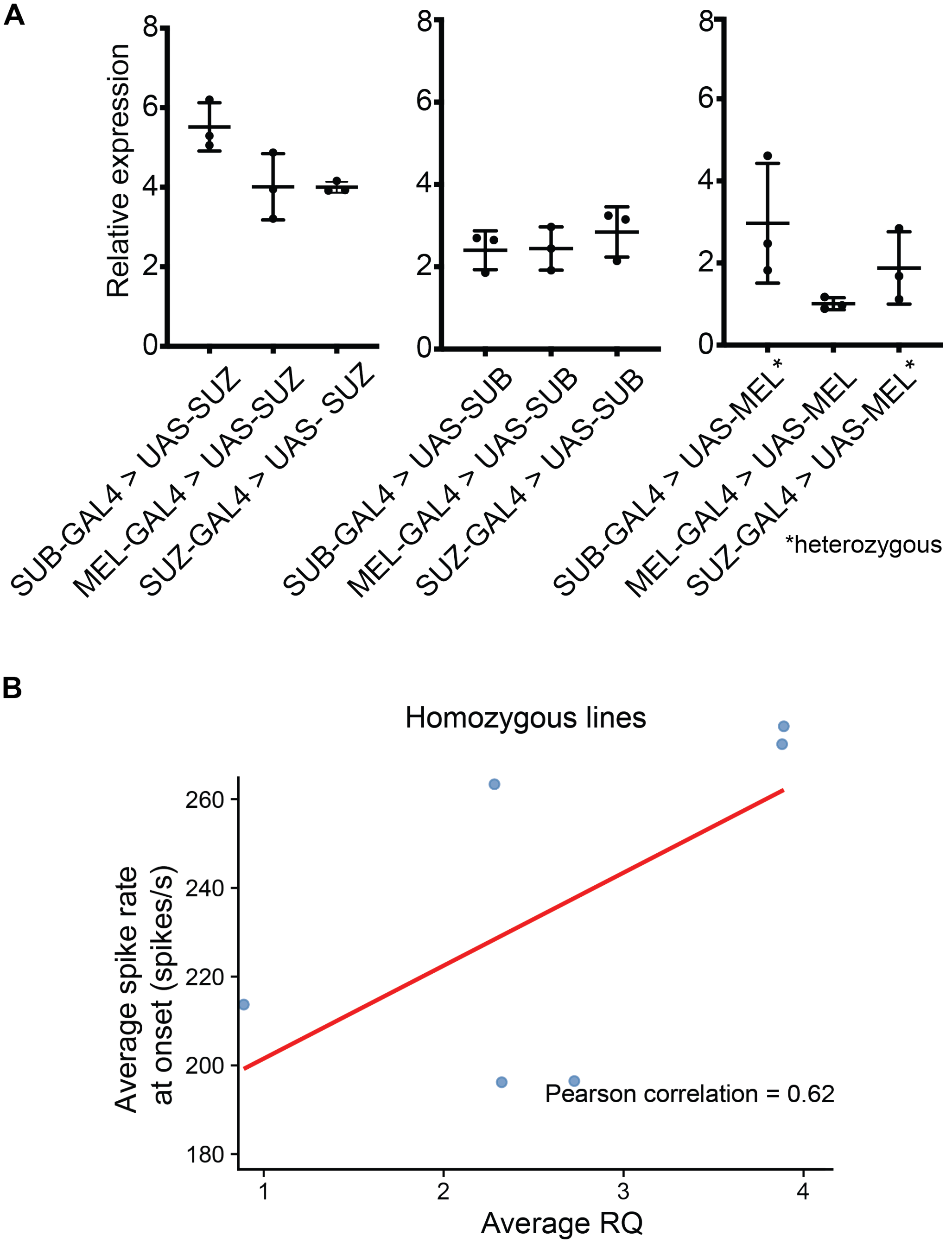


S4 Fig. Quantitative RT-PCR of the transgenic lines and their correlation with average spike rates. A) qRT-PCR of the transgenic lines, normalized to the line containing homozygote MEL-GAL4 and UAS-MEL. Note that we were not able to obtain homozygous flies for two strains at the time of the experiment. B) Correlation between average spike rate and expression levels for the six lines that have homozygous flies for both quantitative RT-PCR and SSE recordings.
